# FAIM opposes stress-induced loss of viability and blocks the formation of protein aggregates

**DOI:** 10.1101/569988

**Authors:** Hiroaki Kaku, Thomas L. Rothstein

**Affiliations:** Center for Immunobiology Western Michigan University Homer Stryker MD School of Medicine, 1000 Oakland Drive, Kalamazoo, MI 49008.

## Abstract

A number of proteinopathies are associated with accumulation of misfolded proteins, which form pathological insoluble deposits. It is hypothesized that molecules capable of blocking formation of such protein aggregates might avert disease onset or delay disease progression. Here we report that Fas Apoptosis Inhibitory Molecule (FAIM) counteracts stress-induced loss of viability. We found that levels of ubiquitinated protein aggregates produced by cellular stress are much greater in FAIM-deficient cells and tissues. Moreover, in an in vitro cell-free system, FAIM specifically and directly prevents pathological protein aggregates without participation by other cellular elements, in particular the proteasomal and autophagic systems. Although this activity is similar to the function of heat shock proteins (HSPs), FAIM, which is highly conserved throughout evolution, bears no homology to any other protein, including HSPs. These results identify a new actor that protects cells against stress-induced loss of viability by preventing protein aggregates.

## Introduction

A number of proteinopathies such as neurodegenerative diseases are associated with accumulation of damaged, misfolded proteins that form pathological insoluble deposits, including Alzheimer’s disease (AD), Parkinson’s disease (PD), Huntington’s disease (HD), amyotrophic lateral sclerosis (ALS), and, possibly, chronic traumatic encephalopathy^1,2^. It is hypothesized that molecules capable of blocking aggregation of disease-associated proteins might prevent the onset, or delay progression, of a number of neurodegenerative illnesses^3–6^. Here we report a new mammalian protein that is active against protein aggregates, Fas Apoptosis Inhibitory Molecule (FAIM, also termed FAIM1).

FAIM was originally cloned as a FAS antagonist in mouse primary B lymphocytes^7^. A subsequent study identified the alternatively spliced form, termed FAIM-Long (L)^8^, which has 22 additional amino acids at the N-terminus. Thus, the originally identified FAIM was renamed FAIM-Short (S)^8^. FAIM-L is expressed almost exclusively in the brain and in the testis whereas FAIM-S is ubiquitously expressed^8^. Recently, the *faim-Gm6432* gene, thought to be duplicated from the original *faim* gene, was identified in Muroidea rodents and its expression is limited to the testis^9^.

Intriguingly, *in silico* analysis indicates the existence of *faim* genes in the premetazoan genomes of single-celled choanoflagellates like *M.brevicollis* and *S.rosetta,* which is one of the closest living relatives of animals and a progenitor of metazoan life that first evolved over 600 million years age^10,11^. *S.rosetta* contains only 9411 genes, out of which 2 *faim* genes were found^11^. This evidence suggests that the *faim* gene evolved much earlier than many other genes and domains found in multicellular organisms, including the death domain involved in animal cell apoptosis^12–14^, and implies that this gene may have another major function beyond apoptosis regulation. However, a lack of known consensus effector/binding motifs and even partial sequence homology of FAIM with any other protein has to date rendered it difficult to predict such functions^7^.

A series of overexpression studies demonstrated that FAIM produces resistance to FAS (CD95)-mediated apoptosis in B lymphocytes^7^, HEK293T cells^15^ and PC12 cells^16^, enhances CD40-mediated NF-κB activation in B lymphocytes^17^, and induces neurite outgrowth in the PC12 cell line^18^. Thus, FAIM expresses multiple activities related to cell death, signaling, and neural cell function. Nonetheless, the overarching physiological role of FAIM still remained unclear due to a lack of obvious phenotypic abnormalities of FAIM-deficient mice and cells.

The expression and evolution patterns of the *faim* and *faim-Gm6432* genes suggested that FAIM may be important for testicular functions^9^. Testicular cells are highly susceptible to heat shock^19^ and oxidative stress^20^, which in turn suggested that FAIM might be involved in the cellular stress response. We therefore hypothesized that FAIM might regulate cellular stress response pathways, including the disposition of misfolded proteins that form protein aggregates, in testicular cells or even in other cell types.

This led to the present study, reported here, indicating that FAIM fulfills the previously unknown role of protection against stress in various kinds of cell types. We found FAIM counteracts stress-induced loss of cellular viability. In this process, FAIM binds ubiquitinated proteins and localizes to detergent insoluble material. Importantly, FAIM protects against protein aggregation. These findings strongly suggest a novel, FAIM-specific role in holozoan protein homeostasis.

## Results

### FAIM KO Cells are Susceptible to Heat/Oxidative Stress-Induced Cell Death

To test the activity of FAIM with respect to stress in testicular cells, we established a FAIM-deficient GC-2spd(ts) germ cell line by CRISPR-Cas9 excision and confirmed FAIM-deficiency by western blotting (Fig. S1A). FAIM KO and WT GC-2spd(ts) cells were cultured under stress conditions and cell viability was assayed by 7-AAD staining. After heat shock and after oxidative stress, loss of cell viability in GC-2spd(ts) cells was markedly diminished in the presence of FAIM as compared to its absence (Figs. S1B and S1C). We did not observe a significant difference in FAS-induced cell death between FAIM-sufficient and FAIM-deficient cells (Figs. S1B and S1C).

To exclude the possibility that FAIM protection is limited to germ cells, FAIM-deficient HeLa cells were generated with CRISPR-Cas9 (Fig. S1D). Similar to GC-2spd(ts) cells, FAIM-deficient HeLa cells were highly susceptible to stress-induced cell death (Figs. 1A and 1B), to a greater extent than WT HeLa cells. To confirm these results with high-throughput methodology, we measured supernatant levels of lactate dehydrogenase (LDH) released from dead cells upon stress induction and again we found increased cellular disruption in the face of FAIM deficiency (Figs. 1C and 1D).

**Figure 1.**
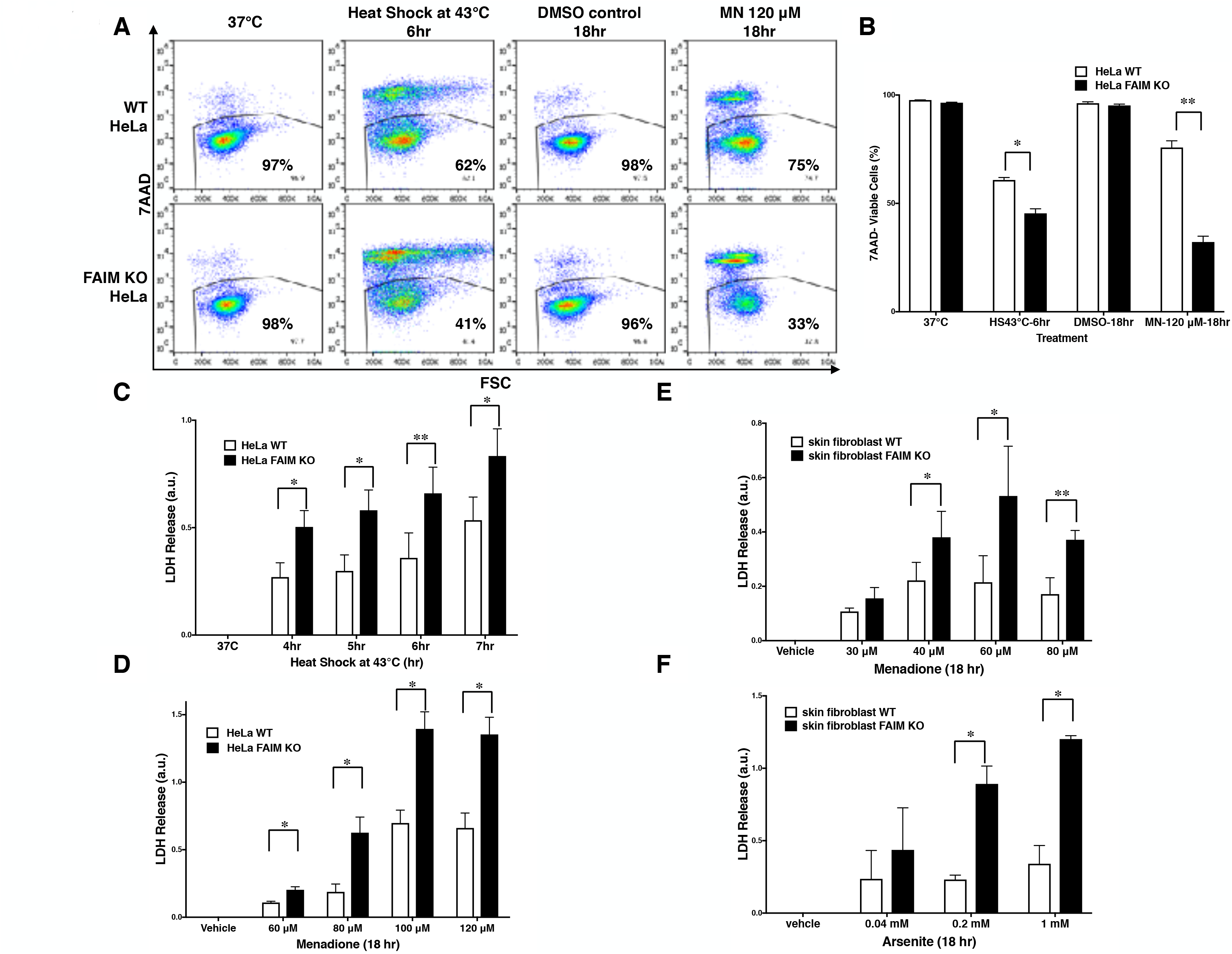
FAIM KO cells are susceptible to heat/oxidative stress-induced cell death. **(A)** WT and FAIM KO HeLa cells were incubated under stress conditions as noted for the indicated periods of time, and then stained with 7-AAD and cell viability was analyzed by flow cytometry. Representative flow data are shown. **(B)** a summary of pooled data from 3 independent experiments (A) are shown. **(C and D)** Cell viability was determined by supernatant LDH leaked from WT and FAIM KO HeLa cells upon heat shock (C) or upon menadione-induced oxidative stress (D), as indicated. Pooled data from 3 independent experiments are shown. (E and F) Primary fibroblasts from WT and FAIM KO mice were subjected to menadione-induced **(E)** and arsenite-induced **(F)** oxidative stress, and cell viability was determined by supernatant LDH. Pooled data from 3 independent experiments are shown. All quantitative data are expressed as mean ± SEM.

To validate the role of FAIM in primary cells, we developed FAIM KO mice in which the mouse *FAIM* gene (Fig. S2A and S2B) was disrupted and we confirmed by WB that FAIM protein was deficient in various tissues from FAIM^−/−^ mice (Fig. S2C). We examined skin-derived fibroblasts which have been shown to be susceptible to menadione-^21^ and arsenite-^22,23^ induced oxidative stress. Consistent with the cell line results, we found vulnerability to oxidative stress induced by menadione (Fig. 1E) and by arsenite (Fig. 1F) to be much greater in FAIM-deficient primary fibroblasts as compared to WT fibroblasts. Thus, data from 3 different cell types indicate that FAIM plays an essential role in protecting cells from heat and oxidative insults.

### Caspase-Dependent Apoptosis and ROS Production are Normal in FAIM KO Cells under Stress Conditions

Oxidative stress and heat shock induce caspase-dependent apoptosis via ROS generation^24^, which could play a role in stress-induced cell death that is affected by FAIM. To address this issue, we first evaluated ROS generation in FAIM-deficient and WT HeLa cells during oxidative stress, using the CellRox deep red staining reagent. We found no difference in stress-induced ROS, regardless of the presence or absence of FAIM (Fig. S3A). We then evaluated caspase activation under stress conditions, using the CellEvent caspase 3/7 detection reagent. We found that caspase 3/7 activity was not increased in FAIM-deficient HeLa cells (Fig. S3B).

To further evaluate stress-induced cell death, we separated cell death into caspase-dependent and caspase-independent forms. We pretreated cells with the pan-caspase inhibitor, Z-VAD-fmk peptide, before adding menadione, and then measured LDH release (Fig. S3C). Z-VAD-fmk has been reported to partially block menadione-induced cell death^25^. We found menadione-induced LDH release was reduced to a small extent in both FAIM-deficient and FAIM-sufficient cells, resulting in similar levels of caspase-dependent apoptosis (Fig. S3D). Importantly, the increased LDH release induced by menadione in FAIM-deficient cells was for the most part resistant to caspase inhibition (Fig. S3E). Thus, menadione-induced cellular dysfunction, which is greatly magnified in the absence of FAIM, is largely caspase independent. In sum, there is no evidence that ROS/caspase-dependent apoptosis plays any role in the improved cellular viability produced by FAIM in the face of stress conditions.

### FAIM Binds Ubiquitinated Proteins after Cellular Stress Induction

Stress-induced cellular dysfunction is often associated with the appearance of disordered and dysfunctional proteins that must be disposed of to maintain cellular viability and stress-induced disordered proteins are tagged with ubiquitin for intracellular degradation via the proteasome system and the autophagic pathway^26,27^. To determine whether stress-induced loss of viability is associated with FAIM binding to ubiquitinated proteins, we again examined FAIM KO HeLa cells. FAIM KO HeLa cells were transfected with FLAG-tagged FAIM proteins and subjected to oxidative stress followed by anti-FLAG IP and western blotting for ubiquitin (Fig. 2A). Separately, FAIM KO HeLa cells were subjected to heat shock and oxidative stress followed by PLA to detect close proximity of FAIM and ubiquitin (Fig. 2B). Both Co-IP and PLA approaches demonstrated stress-induced interaction between FAIM and ubiquitinated protein. These data indicate that FAIM and ubiquitinated proteins associate with each other in response to cellular stress induction that, as noted above, is less deleterious to FAIM-sufficient HeLa cells as opposed to FAIM-deficient HeLa cells.

**Figure 2.**
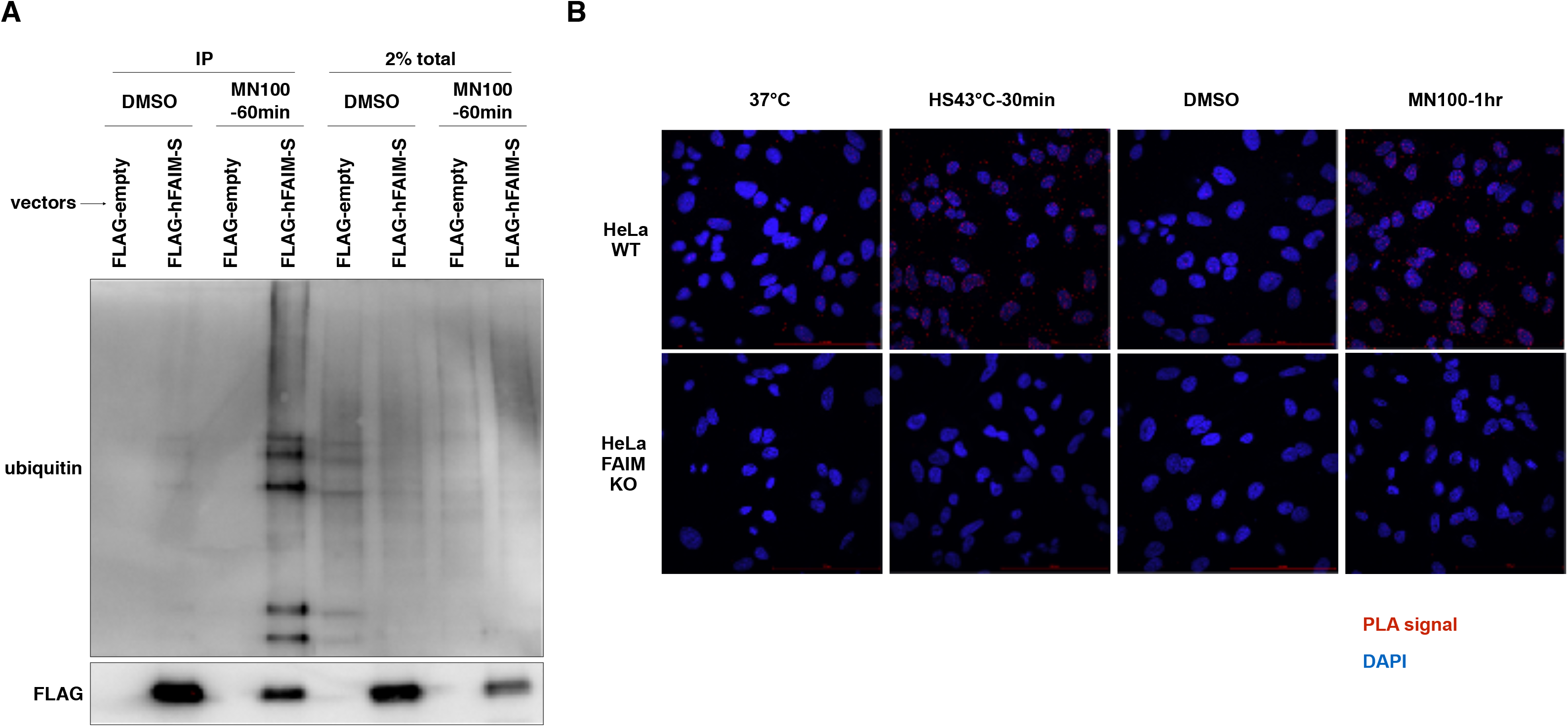
FAIM Binds Ubiquitinated Proteins after Cellular Stress Induction. FAIM-ubiquitin binding was assessed by co-immunoprecipitation **(A)** and *in situ* PLA **(B)**. (A) FAIM KO HeLa cells were transiently transfected with FLAG-tagged FAIM protein. Transfected FAIM KO HeLa cells were subjected to oxidative stress by incubation with menadione (MN, 100 μM) for 1 hour, or were incubated with DMSO (the diluent for menadione), after which cells were harvested. Lysates were immunoprecipitated with anti-FLAG and subjected to SDS-PAGE and western blotted for ubiquitin. (B) FAIM KO HeLa cells and WT HeLa cells were subjected to heat shock (HS) at 43 ^o^C or oxidative stress (MN, 100 μM), as indicated, and then fixed and permeabilized, after which PLA reaction was carried out to detect proximity of FAIM and ubiquitin. Red dots indicate PLA positive signals and nuclei are stained blue with DAPI. Similar results were obtained from at least 2 independent experiments.

### FAIM-Deficient Cells Accumulate Ubiquitinated, Aggregated Proteins in the Detergent-Insoluble Fraction after Stress Induction

The associations among FAIM, ubiquitinated proteins and detergent insoluble material, induced by stress (Fig. 2) suggest that impaired viability in stressed FAIM-deficient cells may be due to accumulation of cytotoxic, ubiquitinated protein aggregates that have overwhelmed disposal mechanisms. To determine if ubiquitinated protein aggregates increase after stress and do so disproportionately in the absence of FAIM, we assessed stress-induced accumulation of ubiquitinated proteins in FAIM KO HeLa cells vs WT HeLa cells by western blotting. We found ubiquitinated proteins accumulated in detergent-insoluble fractions after heat shock (Fig. 3A) and after oxidative stress (Fig. 3B) and did so to a much greater extent in FAIM KO HeLa cells compared to WT HeLa cells (Figs. 3A and 3B).

**Figure 3.**
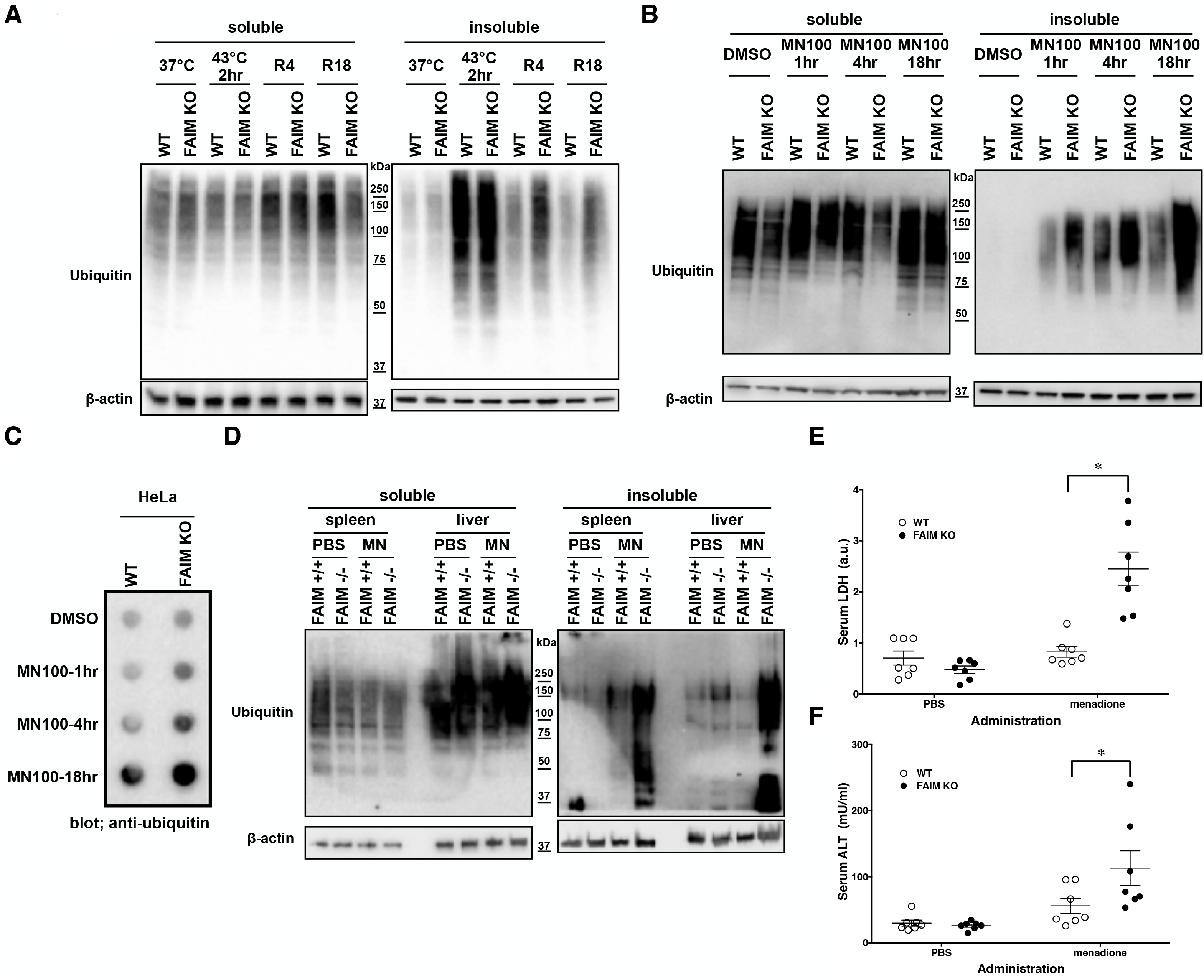
FAIM-deficient cells/tissues accumulate ubiquitinated, aggregated proteins in the detergent-insoluble fraction after stress induction. **(A)** WT HeLa cells and FAIM KO HeLa cells were incubated at 37°C or subjected to heat shock at 43°C for 2 hours followed by recovery at 37°C for 4 hours (R4) and for 18 hours (R18). Cells were lysed and detergent soluble and detergent insoluble fractions were isolated. Equal amounts of protein for each fraction were analysed by western blotting for ubiquitin and actin as a loading control. **(B)** WT HeLa cells and FAIM KO HeLa cells were incubated with menadione (MN) at 100 μM for the times indicated, or were incubated with DMSO vehicle. Cells were then handled as in (A). **(C)** WT HeLa cells and FAIM KO HeLa cells were incubated with menadione (MN) at 100 μM for the indicated times, after which aggregated proteins were filter trapped and blotted with anti-ubiquitin. **(D)** Spleen and liver tissue from FAIM KO mice and their littermate controls were collected 18 hours after intraperitoneal administration of PBS or menadione (MN, 200 mg/kg). Tissue lysates were immediately extracted and protein samples were subjected to SDS-PAGE and western blotted for ubiquitin and actin as a loading control. Results shown in (A, B, C and D) are representative of at least 3 independent experiments. (E and F) Serum samples obtained from mice treated as in (D) were analyzed for content of LDH **(E)** and ALT **(F)**. Data represent mean ± SEM (n=7).

Next, to confirm that ubiquitinated proteins detected by western blotting in the detergent-insoluble fractions represent aggregated proteins, we performed filter trap assay (FTA) using total cell lysates from cells after oxidative stress. In this assay, large aggregated proteins are not able to pass though the 0.2 μm pore-sized filter and remain on the filter^28^. We observed that more aggregated proteins from FAIM-deficient HeLa cell lysates (Fig. 3C) were trapped on the membrane during oxidative stress as compared to WT HeLa lysates. The same was true for primary mouse skin-derived fibroblasts from FAIM KO mice as compared to fibroblasts from WT mice (Figs. S4A and S4B). Thus, following stress, FAIM directly binds ubiquitinated proteins that, when excessive, accumulate in detergent insoluble material, and accumulation of insoluble ubiquitinated proteins is much greater in the absence of FAIM. These results strongly suggest that FAIM is involved in the disposition of stress-induced aggregated proteins, and operates to divert such proteins from a pathway leading to insoluble protein aggregates.

### Ubiquitinated Protein Aggregates Accumulate in FAIM-Deficient Tissues Following Oxidative Stress *in vivo*

To demonstrate that FAIM-deficiency correlates with more insoluble, ubiquitinated protein upon cellular stress *in vivo*, we injected mice with menadione intraperitoneally, and assessed resultant tissue injury^29,30^. Liver and spleen samples were collected 18 hours after menadione administration into FAIM-deficient and littermate control FAIM-sufficient mice, and detergent-soluble and detergent-insoluble proteins were extracted. Similar to our *in vitro* experiments using HeLa cells and primary mouse fibroblasts, in vivo oxidative stress induced dramatically more ubiquitinated proteins in detergent-insoluble fractions from FAIM-deficient liver and spleen cells, as compared to liver and spleen cells from menadione-treated WT mice (Fig. 3D). In accordance with these results, we found much higher levels of menadione-induced serum LDH (Fig. 3E) and ALT (Fig. 3F), which are signs of cell injury and death, in FAIM KO as compared to WT mice. These data indicate that FAIM plays a non-redundant role in preventing accumulation of insoluble, ubiquitinated, aggregated protein in stress-induced animals, and in protecting against cell death.

### FAIM KO Cells Accumulate Aggregation-Prone Proteins

To directly assess the role FAIM plays in prevention of protein aggregation, we employed pulse shape analysis (PulSA)^31^ by flow cytometry to determine the level of aggregated protein. We transiently transfected HeLa cells with eGFP-tagged aggregation-prone proteins (mutant huntingtin exon1) (Figs. 4A), which spontaneously form aggregates in some cells. We found that cells containing aggregated proteins had a narrower and higher pulse shape of eGFP fluorescence than those that did not express protein aggregates, as previously reported (Figs. 4B)^31^. Regardless of transfection efficiency (Figs. 4A), the fraction of HeLa cells expressing aggregated proteins (mutant huntingtin exon 1, Fig. 4B) was significantly higher in FAIM-deficient HeLa cells than in WT HeLa cells, especially at late stages after transfection (Figs. 4C). These results were further verified by FTA. We found that much more aggregated mutant huntingtin (Fig. 4D) was filter trapped in FAIM-deficient HeLa cells than in WT HeLa cells. These results demonstrate the essential role of FAIM in altering the fate of mutant aggregation-prone proteins that play a role in some forms of neurodegenerative diseases.

**Figure 4.**
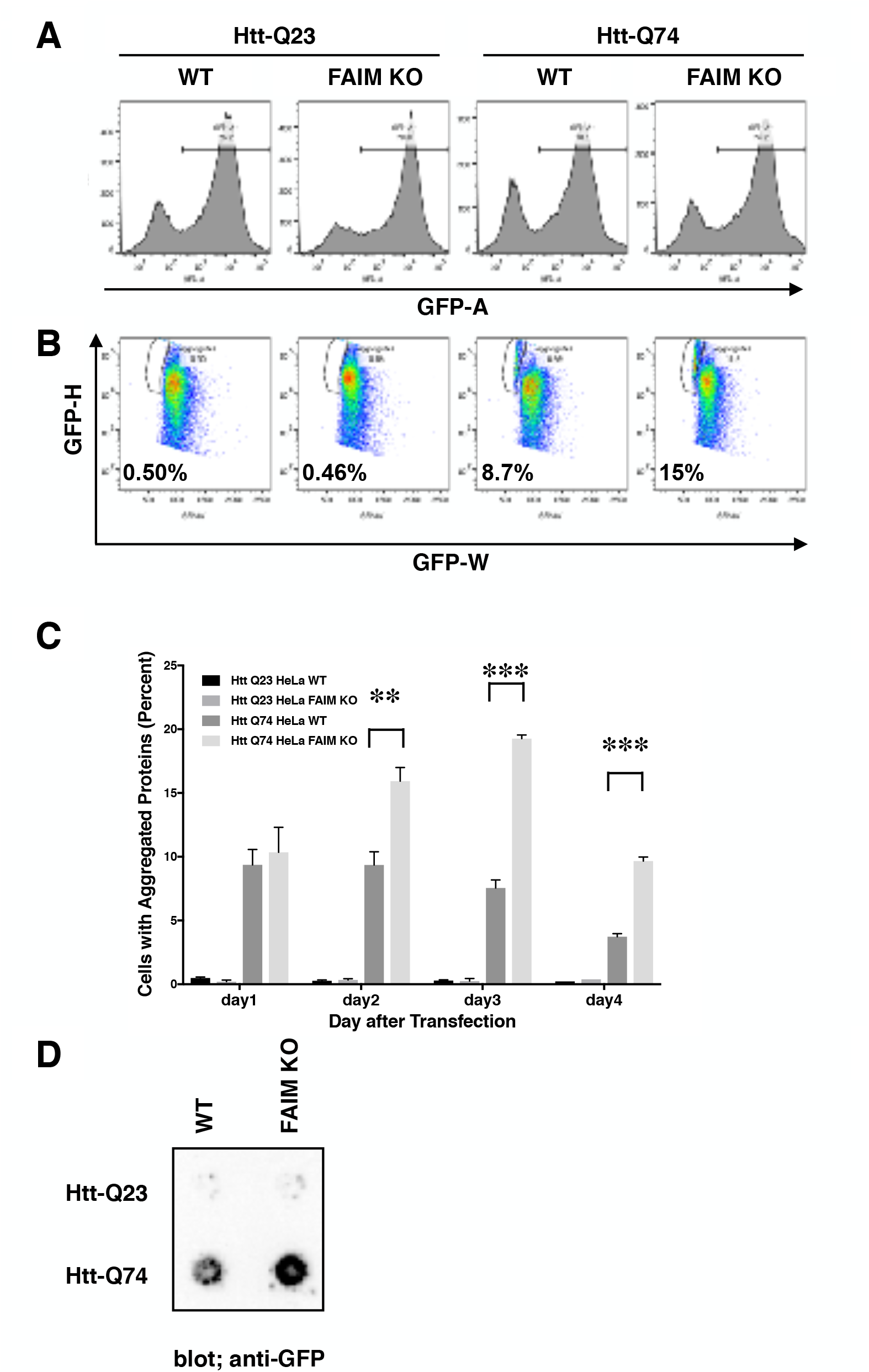
FAIM KO cells accumulate aggregation-prone proteins. **(A)** WT HeLa cells and FAIM KO HeLa cells were transiently transfected with expression vectors for huntingtin (Htt)-Q23 and mutant Htt-Q74 that incorporate an eGFP tag. **(B)** Two days later, eGFP^+^ cells in (A) were evaluated for pulse-width vs pulse-height. The gated area represents cells expressing aggregated proteins. Results representative of 3 independent experiments are shown. **(C)** WT and FAIM KO HeLa cells were transfected as in (A) and harvested at the indicated times. Percentages of cells expressing aggregated proteins out of total eGFP^+^ cells are shown. Data represent mean ± SEM from 3 independent experiments. **(D)** WT and FAIM KO HeLa cells were transfected as in (A) and harvested 2 days later. Equal amounts of total cell lysates were subjected to Filter Trap Assay (FTA) and stained with anti-GFP. Similar results were obtained in at least 3 independent experiments.

### Recombinant FAIM Suppresses Protein Fibrillization/Aggregation in a Cell-Free System

We sought to gain further insight into the molecular mechanism by which FAIM opposes the accumulation of insoluble, aggregated proteins in the face of cellular stress. In order to examine whether FAIM directly inhibits protein aggregation and whether FAIM protein is sufficient to inhibit protein aggregation by itself, without any other cellular components contributing to proteostasis, including elements of the proteasome and autophagic pathways, we mixed recombinant FAIM protein and aggregation-prone β-amyloid (1-42) monomer in an in vitro cell-free system and monitored aggregation status in real-time by ThT fluorescence intensity. We also tested sHSPs, because sHSPs are known to inhibit β-amyloid fibrillization/aggregation in cell-free systems^32,33^ and because HSP27 translocated to detergent-insoluble material in response to stress. We found that β-amyloid aggregation was abrogated in the presence of recombinant FAIM or sHSPs in a dose-dependent manner (Fig. 5A). To confirm these results, aggregation status was assessed by western blotting SDS-PAGE, since aggregated proteins are SDS-resistant. We observed aggregated β-amyloid in the high molecular weight range of negative controls (no added protein control, BSA control). We found the formation of high molecular weight aggregates were dramatically reduced in the presence of recombinant FAIM or sHSPs (Fig. 5B). In addition to β-amyloid, FAIM also inhibited DTT-induced aggregation of α-synuclein A53T mutant protein (Fig. 5C). These data using pure recombinant proteins in a cell-free system indicate that FAIM directly prevents protein fibrillization/aggregation, suggesting that FAIM does not act via known cellular degradation pathways, such as proteasomal and autophagic activities.

**Figure 5.**
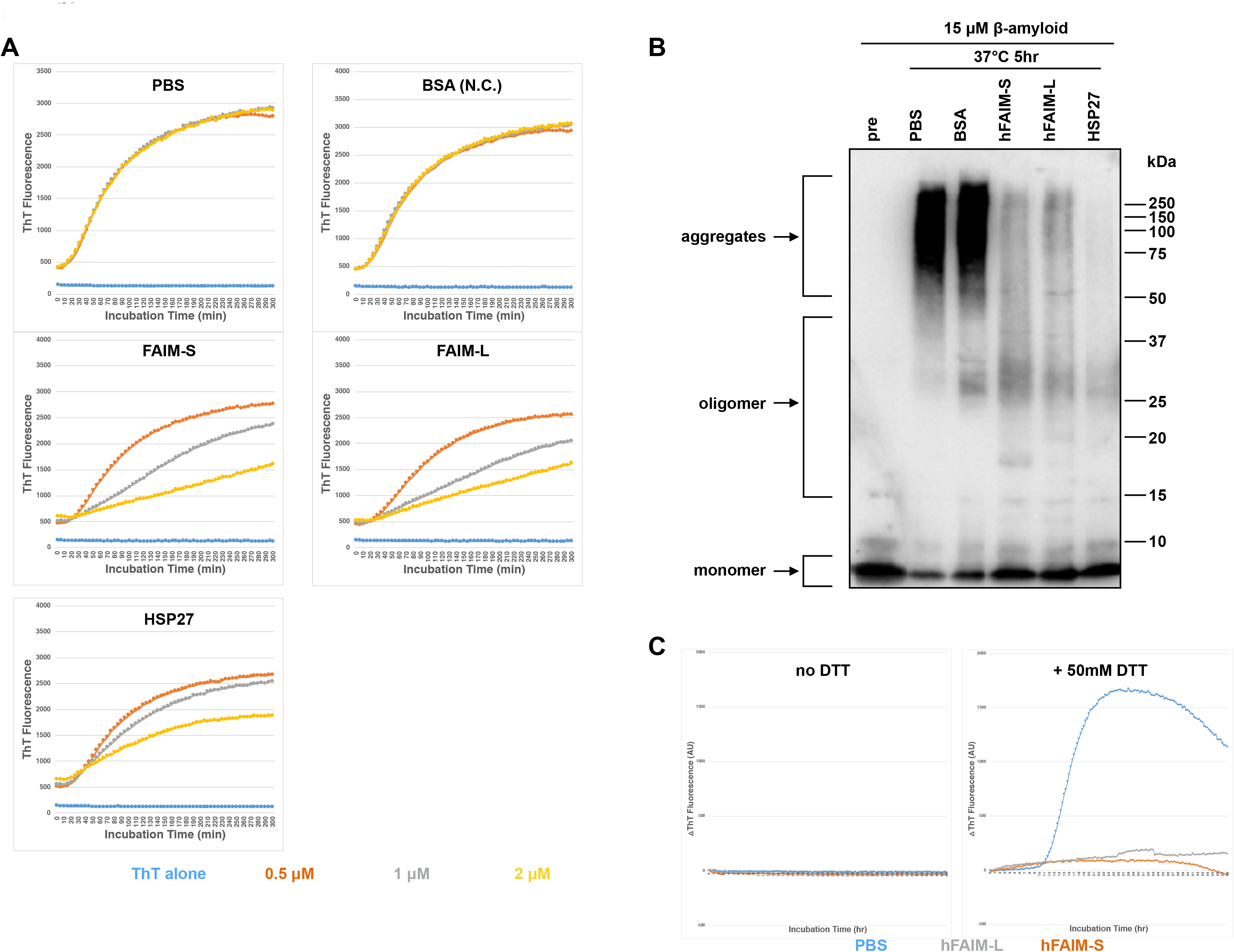
Recombinant FAIM suppresses protein fibrillization/aggregation in a cell-free system. **(A)** Spontaneous aggregation of β-amyloid (15 μM) in vitro was monitored by ThT assay over a period of 5 hours in the presence of recombinant FAIM-S, FAIM-L, HSP27, αB-crystallin or BSA at the doses indicated (blue, ThT alone; orange, 0.5 μM; gray 1 μM; yellow, 2 μM). ThT fluorescence was recorded every 5 minutes. (B) Samples in A at 5 hours were subjected to SDS-PAGE and western blotted for β-amyloid. (C) Aggregation of α-synuclein A53T mutant protein (20 μM) induced by 50 mM DTT (no DTT, left hand panel; 50 mM DTT, right hand panel) was monitored by ThT assay over a period of 48 hours in the presence of recombinant human FAIM-S (5 μM) and FAIM-L (5 μM). ThT fluorescence was recorded every 20 minutes (blue; PBS, orange; FAIM-S, gray; FAIM-L). Representative data from at least 3 independent experiments are shown.

## Discussion

Although the FAIM gene arose in the genomes of the last common holozoan ancestor with a high level of homology among holozoan species, similar to house-keeping genes, its physiological function has been a long-standing enigma. Here, we have demonstrated that FAIM, originally thought of as a FAS-apoptosis inhibitor, plays an unexpected, non-redundant role in protection from cellular stress and tissue damage, leading to improved cellular viability (Figs. 1 and S1). We have elucidated FAIM’s molecular mechanism and demonstrated that it directly interacts with ubiquitinated protein aggregates (Fig. 2), rather than opposing caspase activity or dampening ROS generation (Fig. S3). We have further documented the capacity of FAIM to prevent protein fibrillization/aggregation and shown that this can occur in the absence of other cellular activities (Fig. 5), strongly suggesting that FAIM plays a distinctive, non-redundant, protective role in advanced organisms, for which no other presently known metazoan protein can fully substitute.

Cells and tissues are continuously subjected to environmental insults such as heat shock and oxidative stress, which cause accumulation of cytotoxic, aggregated proteins, leading to caspase-independent cell death. Organisms have evolved protective cellular mechanisms such as HSPs in order to prevent and counteract tissue and organ damage. Our data support the role of FAIM as a new player that antagonizes cytotoxic protein aggregates formation and is not complemented by HSPs.

FAIM expression, translocation, and function closely resemble those of sHSPs in many respects, up to and including prevention of protein aggregation. Although there is no homology between FAIM and sHSPs, implying they have evolved independently, we found structural and biochemical similarities that suggest they contribute to some related common functions in cells. First, the predicted molecular weight of monomer FAIM ranges from 17-43 kDa throughout the evolution of holozoans while sHSPs have a similar monomeric molecular weight range of 15-40 kDa^34^. Second, three-dimensional protein analysis by NMR reveals that FAIM contains a rare compact non-interleaved seven stranded β-sandwich structure with an anti-parallel β-sheet in the C-terminal region and a highly disordered N-terminal region, and experiences temperature-dependent conformational changes^15,35^. Highly disordered protein regions have been suggested to be important in binding aggregation-prone targets^36^. Strikingly, similar structural features are observed in sHSPs^37,38^. Third, amino acid composition analysis of FAIM protein sequences from choanoflagellate to human shows significant underrepresentation of cysteine residues (only 1.4%), which is also true of sHSPs^39^. A reduced number of cysteines may be important to prevent unwanted crosslinking under oxidative conditions, enabling FAIM and sHSPs to resist denaturation and perform their common function to prevent protein aggregation induced by cellular oxidative stress^39^.

Additional evidence supporting similar functions of FAIM and sHSPs was obtained from overexpression experiments. First, NGF-induced neurite outgrowth in vitro was promoted by overexpression of FAIM in the PC12 cell line^18^ and by overexpression of HSP27 in dorsal root ganglion neurons^40^. Second, FAS-mediated apoptosis was inhibited by overexpression of FAIM in B lymphocytes^7^, in PC12 cells^16^ and in HEK293T cells and cortical neurons^15^, and by overexpression of HSP27 in L929 cells^41^. Finally, NF-κB activation was enhanced by overexpression of FAIM in NGF-treated PC12 cells^18^ and in CD40-stimulated B lymphocytes^17^ whereas overexpression of HSP27 enhanced NF-κB activation in TNFα-treated U937 cells and MEF cells^42^. However, despite these many similarities between FAIM and sHSPs, FAIM is not a sHSP. FAIM is not homologous with sHSPs and contains no α-crystallin domain, and, most importantly, the function of FAIM is not complemented by sHSPs in FAIM-deficient cell lines, primary cells, or animals.

The aggregation of proteins into fibrillar high molecular-weight species is a hallmark of numerous human neurodegenerative disorders^43^. In the situation where overwhelming generation of misfolded or aggregated proteins due to cellular or aging stress occurs, these cytotoxic species must be degraded. However, in normal aged neuronal cells, autophagy-related genes are downregulated, leading to dysfunction of autophagy-mediated aggregate clearance^44^. Interestingly, autophagy was found to be impaired in HD model mice and HD patients^45^. Further, proteasomal function has been reported to decline with age^46^. Thus, in a situation of low autophagic and proteasomal activity, the role of FAIM in preventing aggregation may be especially critical to maintaining proteostasis.

The pathophysiology of the neurodegenerative disorder, AD, involves protein aggregates in the form of β-amyloid plaques and tau neurofibrillary tangles^6^. An association between AD and FAIM has been suggested^47^. FAIM-L expression was found to be impaired in the brains of AD patients, especially in late BRAAK stages^47^. Given that FAIM protein prevented β-amyloid aggregation in vitro (Figs. 5A and B), we suggest the novel hypothesis that low/no FAIM expression might be pathogenically linked to more rapid, aggressive, overwhelming β-amyloid aggregation in AD patients rather than simply functioning as a marker of AD progression. Consistent with this hypothesis, we show in this study that oxidative stress, applied in vivo to intact animals, produces increased levels of protein aggregates in the face of FAIM deficiency (Figs. 3 and 4). Accumulated data suggest that oxidative stress is linked to neurodegenerative diseases, presumably due to accumulation of toxic protein aggregates triggered by oxidative stress-induced protein oxidation^48–50^. Thus, diminished FAIM expression in AD patients, combined with our finding of increased protein aggregation in several organs after oxidative stress delivered to FAIM-deficient mice, suggests that FAIM is pathophysiologically relevant to current paradigms for the origin and/or progression of neurodegenerative diseases that involve disruption of proteostasis. Taken together, our work provides new insights into the interrelationships among FAIM, protein aggregation and cell viability that may have applicability to neurodegenerative diseases.

## Methods

### Ethic Statement

All the mouse studies were performed at the Western Michigan University Homer Stryker MD School of Medicine campus in accordance with guidelines and protocols approved by IACUC committee (IACUC Protocols No. 2016-006 and No. 2017-007).

### Antibodies

Rabbit anti-GFP, rabbit anti-ubiquitin, mouse anti-α-tubulin, rabbit anti-β-amyloid, goat anti-rabbit IgG-HRP-linked and horse anti-mouse IgG-HRP-linked antibodies were obtained from Cell Signaling Technology. Mouse anti-FLAG (M2) antibody and mouse anti-β-actin antibody were obtained from MilliporeSigma. Mouse anti-ubiquitin (UB-1) was obtained from Abcam. Rabbit anti-α-synuclein antibody was obtained from Thermo Fisher Scientific. Affinity purified anti-FAIM antibody was obtained from rabbits immunized with CYIKAVSSRKRKEGIIHTLI peptide (located near the C-terminal region of FAIM) as previously described^17^.

### Plasmids

FLAG-tag-hFAIM-S expression vector was constructed using pCMV-(DYKDDDDK)-C (Takara) (cloned into EcoRⅠ and KpnⅠ sites). Primers used for the cloning are shown in Table S1. The insert was verified by sequencing (Genewiz).

The following plasmids were obtained from Addgene.

EGFP-α-synuclein-A53T^51^, plasmid #40823

pEGFP-Q74^52^, plasmid #40262

pEGFP-Q23^52^, #40261

pSpCas9(BB)-2A-GFP (PX458)^53^, plasmid #48138

### Generation of FAIM-deficient mice

FAIM-deficient (KO) mice were generated in conjunction with the inGenious Targeting Laboratory. The target region, including the *faim* coding regions of exons 3-6 (9.58 kb), was replaced by sequences encoding eGFP and neomycin-resistant genes (Fig. S2A). The targeting construct was electroporated into ES cells derived from C57BL/6 mice. Positive clones were selected by neomycin and screened by PCR and then microinjected into foster C57BL/6 mice. Subsequent breeding with wild-type C57BL/6 mice produced F1 heterozygous pups. Offspring from heterozygous mice were selected using PCR. Mice were maintained on a C57BL/6 background. We performed genotyping PCR using genomic DNA from ear punches with a mixture of 4 primers to identify the wild-type allele and the mutant alleles, generating 514bp and 389bp DNA amplicons, respectively (Fig. S2B). Primers are shown in Table S1. Mice were cared for and handled in accordance with National Institutes of Health and institutional guidelines. FAIM-KO mice were viable, developed normally and did not show any obvious phenotypic changes in steady state conditions (data not shown). The heterozygous intercrosses produced a normal Mendelian ratio of FAIM^+/+^, FAIM^+/−^, and FAIM^−/−^ mice.

### Cell culture and transfection

HeLa, GC-1 spg and GC-2spd(ts) cell lines were obtained from the American Type Culture Collection (ATCC). HeLa cells were cultured in DMEM medium (Corning) whereas GC-2spd(ts) cells were cultured in EMEM (Corning). Both DMEM and EMEM contained 10% FCS, 10 mM HEPES, pH 7.2, 2 mM L-glutamine and 0.1 mg/ml penicillin and streptomycin. Transfection was performed using Lipofectamine 2000 for GC2spd(ts) cells or Lipofectamine 3000 for HeLa cells, according to the manufacturer’s instructions (Themo Fisher Scientific). Primary fibroblasts were purified and cultured as previously described^54^. Briefly, skin in the underarm area (1 cm x 1 cm) was harvested in PBS. The tissue was cut into 1 mm pieces. To extract cells, tissues were incubated at 37 ^o^C with shaking in 0.14 Wunsch units/ml Liberase Blendzyme 3 (MilliporeSigma) and 1 x antibiotic/antimycotic (Thermo Fisher Scientific) in DMEM/F12 medium (Corning) for 30 – 90 min until the medium became ‘’fuzzy’’. Tissues were washed with medium three times and then cultured at 37 ^o^C. After 7 days, cells were cultured in EMEM containing 15% FBS plus penicillin/streptomycin for another 7 days. Cells were used at this point for experiments.

### Generation of FAIM knockout cell lines with CRISPR/Cas9

Guide RNA (gRNA) sequences for both human and mouse FAIM gene were designed using a CRISPR target design tool (http://crispr.mit.edu) in order to target the exon after the start codon^53^. Designed DNA oligo nucleotides are shown in Table S1. Annealed double strand DNAs were ligated into pSpCas9(BB)-2A-GFP (PX458) vector (Addgene) at the Bpi1 (Bbs1) restriction enzyme sites using the ‘Golden Gate’ cloning strategy. The presence of insert was verified by sequencing. Empty vector was used as a negative control. Transfection was performed using lipofection and a week after the transfection, eGFP^+^ cells were sorted with an Influx instrument (Becton Dickinson), and seeded into 96 well plates. FAIM knockout clones were screened by limiting dilution and western blotting.

### In vitro cellular stress induction

To induce mild heat shock, cells in culture dishes were incubated in a water bath at 43 ^o^C for the indicated period^55^. In some experiments, cells were recovered at 37 ^o^C after heat stress induction at 43 ^o^C for 2 hours or more as previously described^55^. To induce oxidative stress, menadione (MN) (MilliporeSigma), dissolved in DMSO at 100 mM, was added to medium at the indicated final concentration for 1 hour. In oxidative stress experiments where cells were harvested at time points beyond 1 hour with menadione, cells were washed once with medium and fresh medium (without menadione) was added to the cell culture as previously described^56^. To induce FAS-mediated apoptosis in GC-2spd(ts), cells were cultured with 5 μg/ml anti-FAS antibody (clone; Jo2, BD Pharmingen) as previously described^57^.

### In vivo mouse stress induction

Acute oxidative stress was induced by a single intraperitoneal injection of menadione (200 mg/kg in PBS) into mice^29,30^, and then mice were euthanized 18 hours after the injection. Spleens and livers were removed and protein was immediately extracted for western blotting analysis. All experimental protocols were approved by the Western Michigan University’s institutional and/or licensing committee.

### Cell viability analysis with flow cytometry

Adherent cells were detached by Trypsin-EDTA. Adherent and floating cells were harvested and pooled, after which cells were resuspended in 2 μg/ml 7-aminoactinomycin D (7-AAD) (Anaspec). Cell viability was assessed using Gallios (Beckman Coulter) or Attune (Thermo Fisher Scientific) flow cytometers. Data were analyzed using FlowJo v9 or v10 software (FlowJo, LLC).

### Viability analysis by released LDH detection

Following stress induction in vitro or in vivo, LDH released into the supernatant, or into the serum, from damaged cells was quantified using the Cytotox 96 Non-radioactive Cytotoxicity Assay (Promega). Serum samples were diluted in PBS (1:20).

### ALT activity assay

Following stress induction in vivo, serum was harvsted and ALT levels were monitored using the ALT Activity Assay Kit (BioVision). OD at 570 nm (colorimetric) was detected with a Synergy Neo2 instrument. Serum samples were diluted in ALT assay buffer (1:5).

### Western blotting

Cells were washed twice with PBS and lysed in RIPA lysis buffer (1% Nonidet P-40, 0.5% sodium deoxycholate, 0.1% SDS, 150 mM NaCl, 50 mM Tris-HCl (pH 8.0), 2 mM EDTA) containing supplements of 2 mM Na_3_VO_4,_ 20 mM NaF, and a protease inhibitor cocktail (Calbiochem) for 30 min on ice. In addition to the above supplements, 10 mM N-ethylmaleimide (NEM) (MilliporeSigma), 50 μM PR-619 (LifeSensors) and 5 μM 1,10-phenanthroline (LifeSensors) were added in the lysis buffer for ubiquitin detection by western blotting. Lysates were clarified by centrifugation at 21,100 x g for 10 min. Supernatants were used as RIPA-soluble fractions. The insoluble-pellets (the RIPA-insoluble fractions) were washed twice with RIPA buffer and proteins were extracted in 8M urea in PBS (pH 8.0). Protein concentrations were determined using the 660nm Protein Assay Reagent (Pierce).

Protein samples in 1 x Laemmli buffer with 2-ME were boiled for 5 min. Equal amounts of protein for each condition were subjected to SDS-PAGE followed by immunoblotting. Signals were visualized using a ChemiDoc Touch Imaging System (Bio-Rad) and Image Lab software (Bio-Rad)

### Immunoprecipitation

Cells expressing FLAG-tag proteins were lysed in 0.4% Nonidet P-40, 150 mM NaCl, 50 mM Tris-HCl (pH 8.0), 2 mM EDTA, 2 mM Na_3_VO_4,_ 50 mM NaF, and protease inhibitor cocktail for 30 min on ice. Lysates were clarified by centrifugation at 21,100 x g for 10 min. Equal amounts of protein for each supernatant were mixed with anti-FLAG M2 Magnetic Beads (MilliporeSigma) and incubated at 4 ^o^C under gentle rotation for 2 hr. Beads were washed with lysis buffer 4 times and FLAG-tag proteins were eluted with 100 μg/ml 3x FLAG peptide (MilliporeSigma) 2 times. Eluates were pooled and western blotting was performed to detect FLAG-FAIM binding proteins.

### Filter trap assay (FTA)

WT and FAIM KO cells were transiently transfected with eGFP-tagged aggregation-prone protein expression vectors (huntingtin), and fluorescently tagged cells were then harvested at 48 hours to detect protein aggregates. WT and FAIM KO cells were incubated with or without menadione then harvested after the indicated period to detect ubiquitinated protein aggregates. Cells were washed with PBS and then lysed in PBS containing 2% SDS, 1 mM MgCl_2,_ protease inhibitor cocktail and 25 unit/ml Benzonase (MilliporeSigma). Protein concentrations were quantified using 660 nm Protein Assay Reagent with Ionic Detergent Compatibility Reagent (IDCR) (Thermo Fisher Scientific). Equal amounts of protein extracts underwent vacuum filtration through a pre-wet 0.2 μm pore size nitrocellulose membrane (GE Healthcare) for the detection of ubiquitinated protein aggregates or a 0.2 μm pore size cellulose acetate membrane (GE Healthcare) for the detection of huntingtin aggregates using a 96 well format Dot-Blot apparatus (Bio-Rad). The membrane was washed twice with 0.1% SDS in PBS and western blotting using anti-ubiquitin or anti-GFP antibody was carried out to detect aggregated proteins.

### Pulse-shape analysis (PulSA)

Cells expressing eGFP-tagged huntingtin proteins were harvested and eGFP expression was analyzed on an LSR Fortessa (BD Pharmingen) flow cytometer for PulSA analysis to detect protein aggregates, as previously described^31^. Data was collected in pulse-area, height and width for each channel. At least 10,000 cells were analyzed.

### in situ proximity ligation assay (PLA)

Cells were cultured for 24 hours on poly-L-lysine coated coverslips (Corning) in 24-well plates. Cells with or without cellular stress induction were fixed and permeabilized with ice-cold 100% methanol. PLA reaction was carried out according to the manufacturer’s instructions using Duolink In Situ Detection Reagents Orange (MilliporeSigma). ProLong Gold Antifade Reagent with DAPI (Cell Signaling Technology) was used to stain nuclei and to prevent fading of fluorescence. Fluorescence signals were visualized with a Nikon A1R^+^ confocal microscope (Nikon).

### His-tag recombinant protein production

His-tag protein expression vectors were constructed using pTrcHis TA vector according to the manufacturer’s instructions. In brief, PCR amplified target genes were TA-cloned into the vector (Themo Fisher Science) and inserted DNA was verified by sequencing (Genewiz). Proteins were expressed in TOP10 competent cells (Themo Fisher Science) and were purified using a Nuvia IMAC Nickel-charged column (Bio-Rad) on an NGC Quest chromatography system (Bio-Rad). Protein purity was verified using TGX Stain-Free gels (Bio-Rad) on ChemiDoc Touch Imaging System and with Image Lab software and each protein was determined to be >90% pure.

### Thioflavin T fluorescence assay

Fibril/aggregation formation of 15 μM β-amyloid (1-42) (Anaspec) or purified 20 μM α-synuclein A53T, assembled in a 384-well plate (Greiner), was assessed by Thioflavin T (ThT) (MilliporeSigma) fluorescence using a Synergy Neo2 Multi-Mode Microplate Reader (Bio-Tek). Reader temperature was set at 37°C with continuous shaking between reads. ThT fluorescence intensity was measured using an excitation wavelength of 440 nm and an emission of 482 nm. PMT gain was set at 75.

Fluorescence measurements were made from the top of the plate, with the top being sealed with an adhesive plate sealer to prevent evaporation.

### Statistical Analysis

All quantitative data are expressed as mean ± SEM. Student’s *t*-test was used for statistical determinations with GraphPad Prism 7 software. Values of *p*<0.05 are considered statistically significant (\**p*<0.05, \*\**p*<0.01 or \*\*\**p*<0.001).

## Acknowledgments

We are grateful to our colleagues for helpful discussions and technical assistance throughout the course of this study. We also thank Dr. Sharon Singh for her critical reading of the manuscript; Mr. Alex Ludlow for genotyping experiments of FAIM-deficient mice.

## Author contributions

H.K. designed and performed research, analyzed and interpreted data, and wrote the manuscript; T.L.R. analyzed and interpreted data, and wrote the manuscript.

