## Supplementary Figures for "FAIM opposes stress-induced loss of viability and blocks the formation of protein aggregates"

**Title**

**Abstract**

A number of proteinopathies are associated with accumulation of misfolded proteins, which form pathological insoluble deposits. It is hypothesized that molecules capable of blocking formation of such protein aggregates might avert disease onset or delay disease progression. Here we report that Fas Apoptosis Inhibitory Molecule (FAIM) counteracts stress-induced loss of viability. We found that levels of ubiquitinated protein aggregates produced by cellular stress are much greater in FAIM-deficient cells and tissues. Moreover, in an in vitro cell-free system, FAIM specifically and directly prevents pathological protein aggregates without participation by other cellular elements, in particular the proteasomal and autophagic systems. Although this activity is similar to the function of heat shock proteins (HSPs), FAIM, which is highly conserved throughout evolution, bears no homology to any other protein, including HSPs. These results identify a new actor that protects cells against stress-induced loss of viability by preventing protein aggregates.

Figure S1

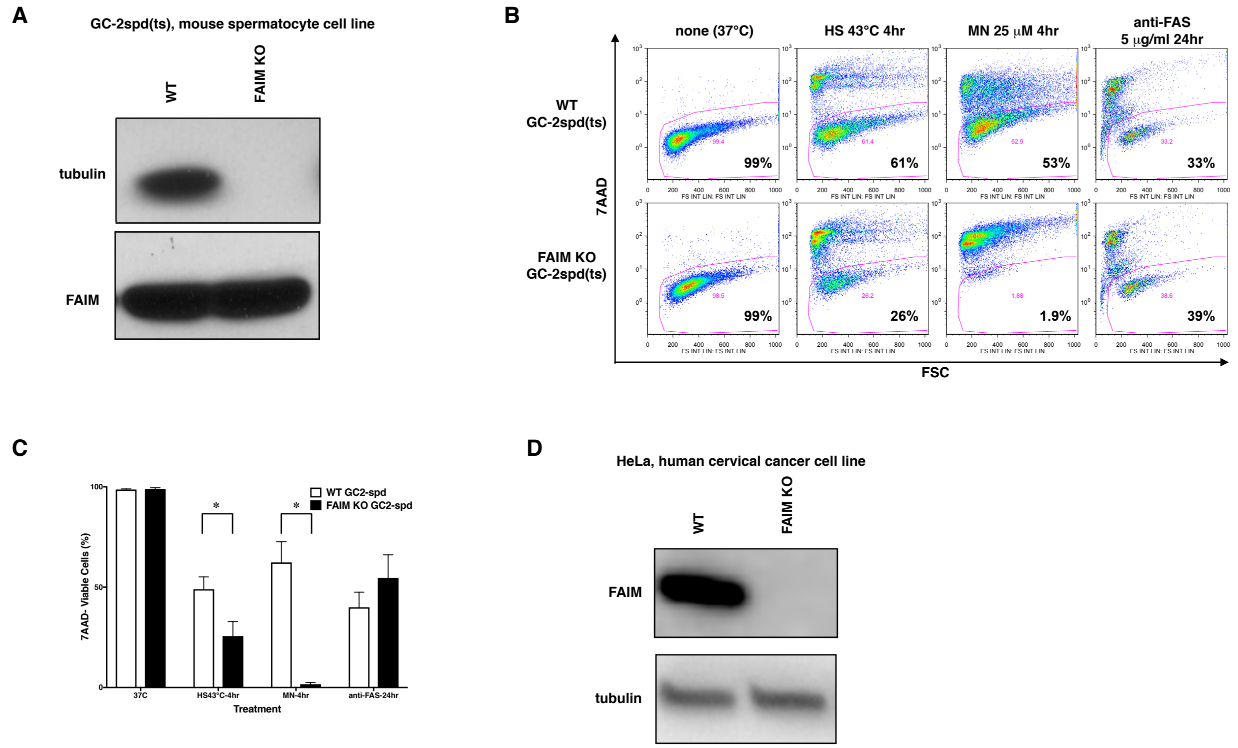


**Figure S1. FAIM KO cells are susceptible to heat/oxidative stress-induced cell death. (A and D)** Western blot analyses of FAIM protein expression levels using cell lysates are shown for GC-2spd(ts) (A) and HeLa cells (D). **(B and C)** GC-2spd(ts) cells, were incubated under stress conditions as noted, for the indicated periods of time. Cell were also exposed to anti-FAS antibody for 24 hours. Cells were stained with 7-AAD and cell viability was analyzed by flow cytometry Representative flow data are shown in (B). A summary of pooled data from 3 independent experiments is shown in **(C)**. Data represent mean ± SEM. HS, heat shock; MN, menadione.

Figure S2

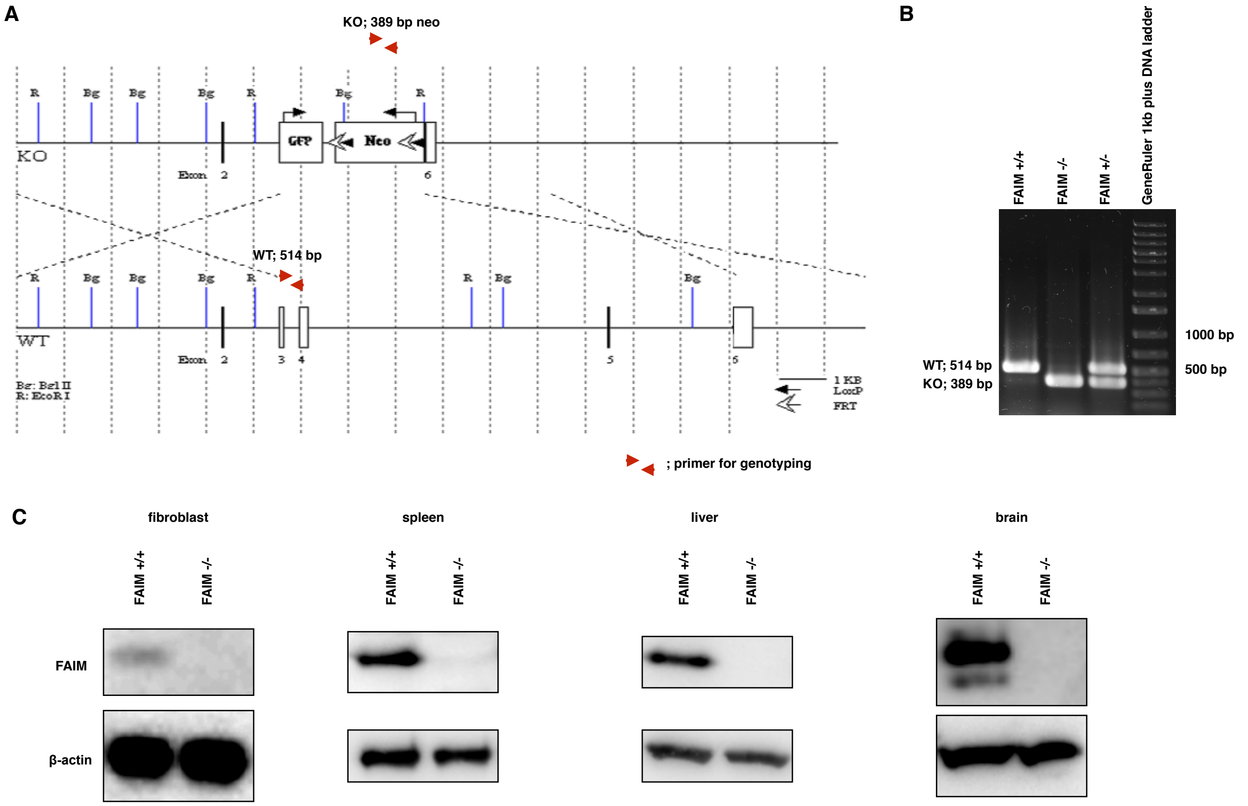


**Figure S2.** ***faim KO m*ice lack exons 3-5**. **(A)** Schematic representation of the targeting vector and the targeted allele of the mouse *faim* gene. **(B)** Genotype determination of *faim* mice by PCR. Multiplex PCR genotyping analyses for KO (389 bp) and WT (514 bp) *faim* genes were performed to confirm the genotypes of wild-type ($^{+/+}$), heterozygous ($^{+/-}$) and homozygous ($^{-/-}$) mice. Representative genotyping results are shown. **(C)** FAIM protein expression was analyzed by western blotting using the indicated tissues from FAIM$^{+/+}$ or FAIM$^{-/-}$mice.

Figure S3

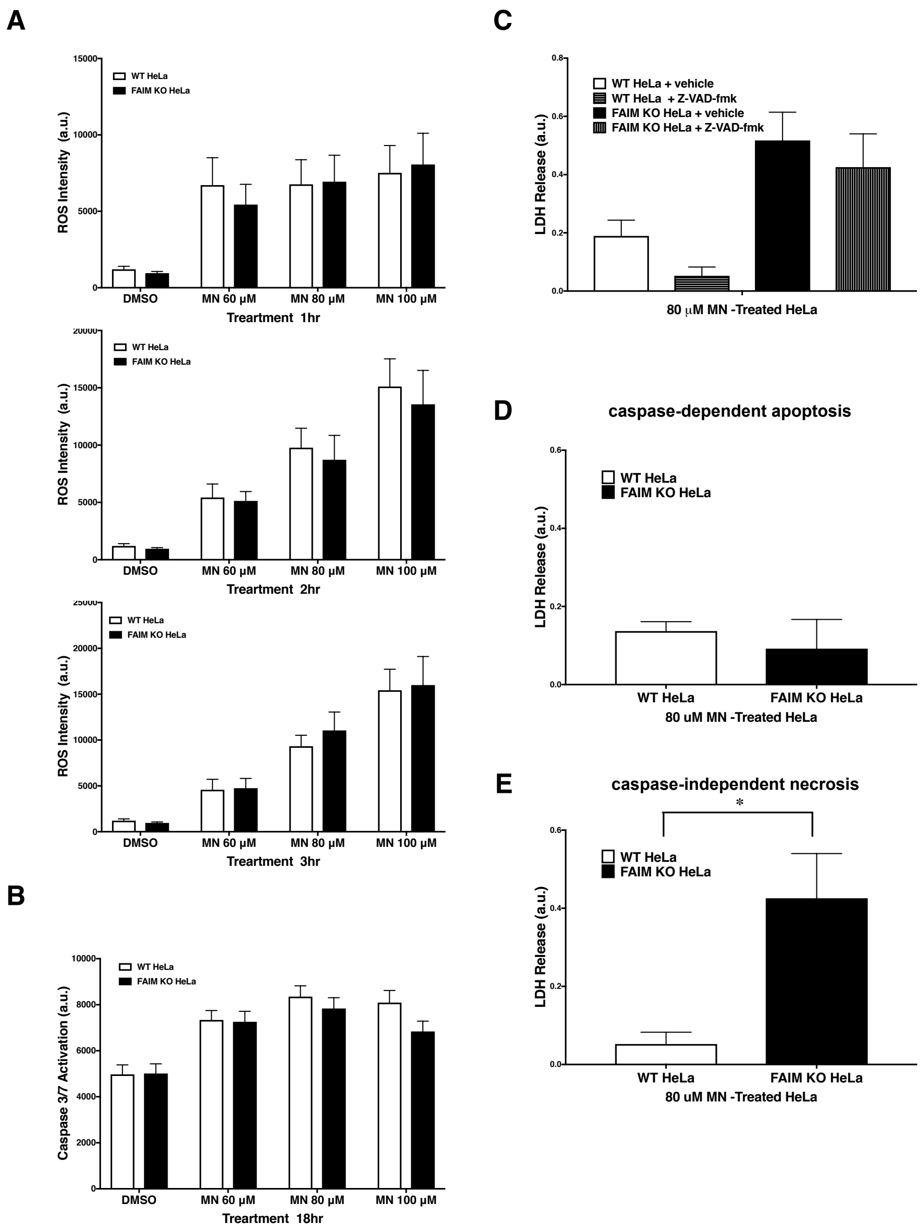


**Figure S3. Caspase-dependent Apoptosis and ROS production are normal in FAIM KO cells under stress conditions. (A)** ROS production was measured by CellRox deep red staining reagent after oxidative stress induction. **(B)** Apoptosis induction was assessed by monitoring Caspase3/7 activation with CellEvent caspase3/7 detection reagent after oxidative stress induction as indicated. **(C, D and E)** Cell disruption was determined by LDH release with or without the pan-caspase inhibitor, Z-VAD-fmk, under oxidative stress conditions as indicated. Caspase-dependent cell death (D) and caspase-independent cell death (E) were calculated based on (C). A summary of pooled data from 3 independent experiments is shown. Data represent mean ± SEM. MN, menadione.

Figure S4

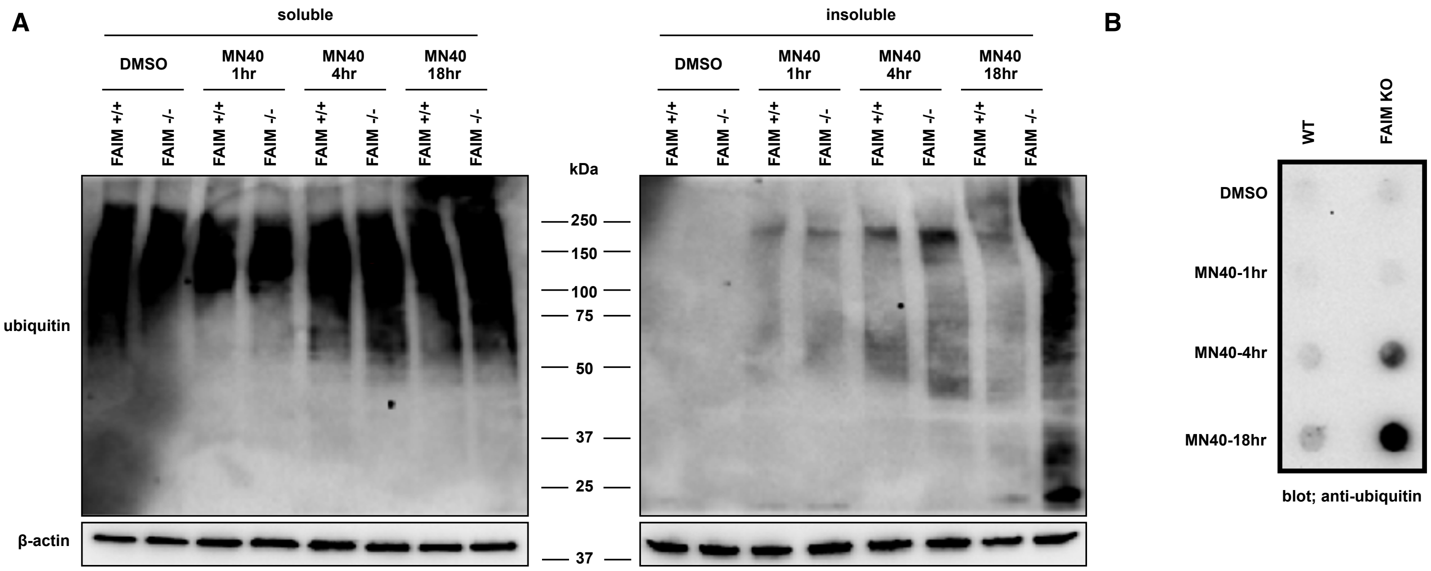


**Figure S4. FAIM-deficient primary fibroblasts accumulate ubiquitinated, aggregated proteins in the detergent-insoluble fraction after stress induction.** **(A)** Primary mouse skin fibroblasts from WT and FAIM KO mice were incubated with menadione (MN) at 40 μM for the times indicated, or were incubated with DMSO vehicle. Cells were lysed and detergent soluble and detergent insoluble fractions were isolated. Equal amounts of protein for each fraction were analysed by western blotting for Ubiquitin, and actin as a loading control. **(B)** Primary mouse skin fibroblasts from WT and FAIM KO mice were incubated with menadione at 40 μM for the times indicated, or were incubated with DMSO vehicle. Aggregated proteins were filter trapped and blotted with anti-ubiquitin. Similar results were obtained from 2 independent experiments and representative data are shown in (A and B).

**Supplemental Materials**

### Supplemental Table 1: Primers/Oligonucleotides Used

| Name | Sequence |
|---|---|
| mouse genotyping for WT allele fwd | ACGGATCTCGTAGCTGTTTGGGACG |
| mouse genotyping for WT allele rev | CCAGCGTGTACTCGTATGCGAAGCC |
| mouse genotyping for FAIM knockout allele fwd | CAGAAGAACTCGTCAAGAAGGC |
| mouse genotyping for FAIM knockout allele rev | CAAGCGAAACATCGCATCGAGCG |
| CRISPR-Cas9 oligo nucleotides for mouse FAIM fwd | CACCGTGACGGATCTCGTAGCTGTTTGG |
| CRISPR-Cas9 oligo nucleotides for mouse FAIM fwd | AAACAACAGCTACGAGATCCGTCAC |
| CRISPR-Cas9 oligo nucleotides for human FAIM fwd | CACCGACAGATCTCGTAGCTGTTTGGG |
| CRISPR-Cas9 oligo nucleotides for human FAIM fwd | AAACAAACAGCTACGAGATCTGTC |
| FAIM-S cloning into pCMV-(DYKDDDDK)-C fwd | ATATAGAATTCTAATGACAGATCTCGTAGCTGTTTGG |
| FAIM-S cloning into pCMV-(DYKDDDDK)-C rev | ATGGTACCACTTGCAATCTCTGGGATTTCT |
| FAIM-S cloning into pTrcHis TA vector fwd | ATGACAGATCTCGTAGCTGTTTGG |
| FAIM-S cloning into pTrcHis TA vector rev | TTAACTTGCAATCTCTGGGATTTC |
| FAIM-L cloning into pTrcHis TA vector fwd | ATGGCATCTGGAGATGACAGTC |
| FAIM-L cloning into pTrcHis TA vector rev | TTAACTTGCAATCTCTGGGATTTC |
| HSP27 cloning into pTrcHis TA vector fwd | ATGACCGAGCGCCGCGTCCCCTT |
| HSP27 cloning into pTrcHis TA vector rev | TTACTTGGCGGCAGTCTCATCGGAT |
| α-synuclein A53T cloning into pTrcHis TA vector fwd | ATGGATGTATTCATGAAAGGACTTTC |
| α-synuclein A53T cloning into pTrcHis TA vector rev | TTAGGCTTCAGGTTCGTAGTCTT |
| sequence primer for pX-458 | TGGACTATCATATGCTTACCGTAACTTGAAAG |
| sequence primer for pTrcHis TA vector | TATGGCTAGCATGACTGGT |
